# Electrokinetic detection of single-molecule phosphorylation

**DOI:** 10.1101/2025.02.24.640009

**Authors:** Quan Wang

## Abstract

Current single-molecule fluorescence experiments lack direct sensitivity to molecular phosphorylation state. We show that electrokinetic properties measured with an anti-Brownian trap resolve the number of phosphorylated sites on individual biomolecules and can monitor phosphorylation cycles with single kinase turn overs. As an application, we directly visualize the rate limiting step in the kinase catalytic cycle.

---

The reversible modification of biomolecules by phosphorylation is a primary mechanism for regulating biological function and thus an observation goal for many biophysical techniques in recent years^1–4^. Here, we develop a method to probe phosphorylation and its dynamics in single-molecule fluorescence experiments. Inspired by analytical charge sensing using capillary electrophoresis^5,6^, our method is based on measuring the mobility difference under a driving electric field, but on individual molecules. Phosphorylation on serine, threonine or tyrosine residues render the biomolecule more negatively charged, thus altering the molecule’s terminal velocity. In a typical experimental scenario (Fig. 1A) with highly negatively charged channel surfaces, weakly-charged biomolecules migrate towards the cathode due to electro-osmosis and phosphorylation decreases the migration velocity (Supplementary Note 1). We integrate mobility sensing with an Anti-Brownian Electrokinetic (ABEL) trap^7,8^, which provides two key advantages. First, the ABEL trap uses electrokinetic flows as the feedback mechanism to counter rapid diffusion, allowing voltage responses of single molecules to be directly characterized^7^. Second, the ABEL trap readily achieves seconds-long observations of individual biomolecules in solution, enabling both high-precision mobility sensing and visualization of dynamics.

**Figure 1.**
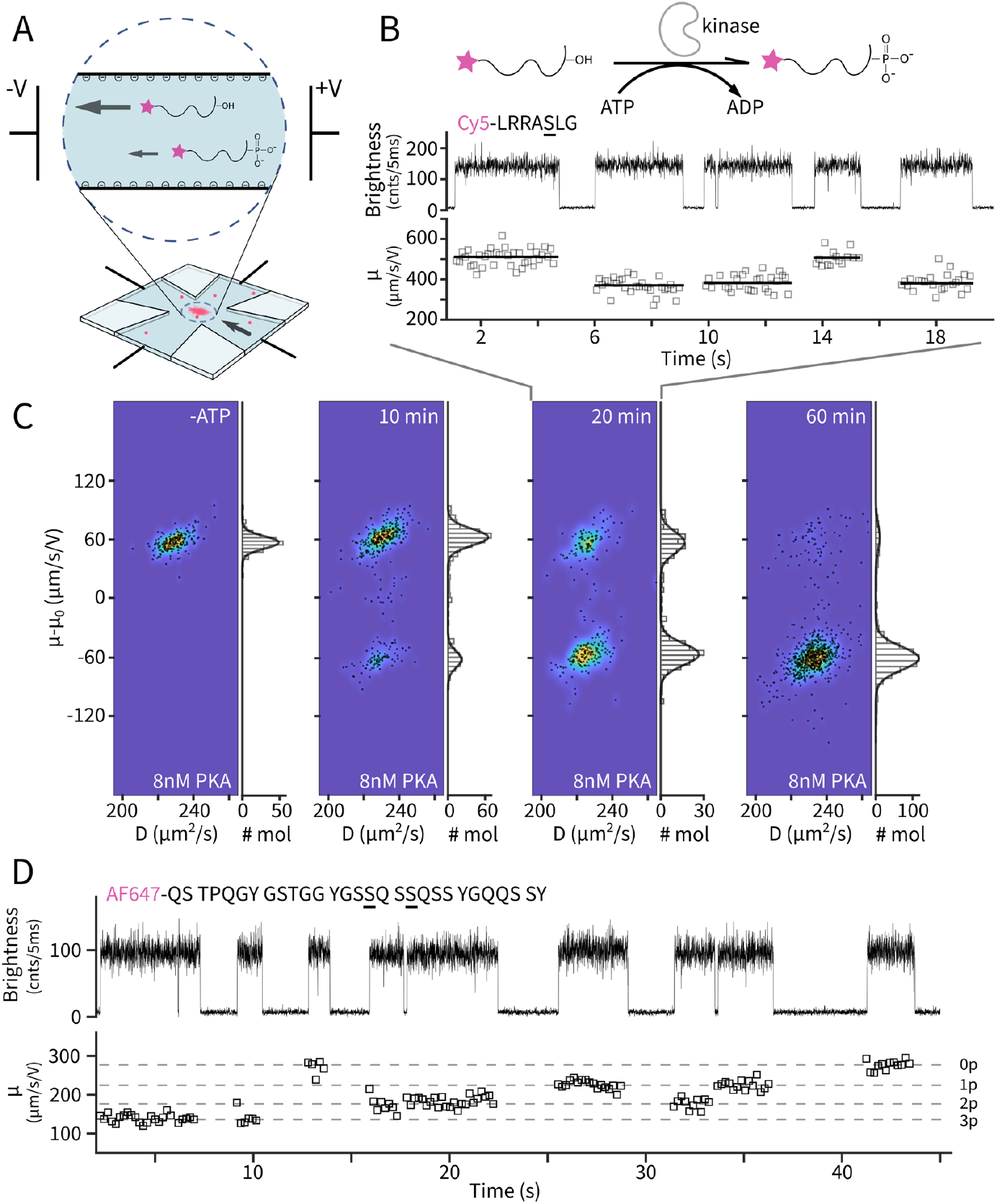
Electrokinetic detection of single-molecule phosphorylation states. **A**. Phosphorylation imparts two negative charges to a single biomolecule of interest, altering its terminal velocity in solution under the applied electric field. This change can be sensed in an ABEL trap (bottom). **B**. Top: reaction scheme and the sequence of the peptide substrate “kemptide” (underscored serine is the phosphorylation site). Bottom: a representative single-molecule time trace of a partial phosphorylation reaction. Brightness is quantified by the number of photons per 5 ms time bins. Electrokinetic mobility is estimated every 100 ms time windows (gray squares). Solid lines represent single-molecule averages. **C**. Diffusivity (*D*)-mobility (*μ*) scatter plots of single molecules in the no-ATP control and the phosphorylation reaction at 10, 20 and 60 min. Pseudo-color represented densities are overlaid. Marginal histograms along *μ* are shown together with Gaussian fits. Mobility values are corrected for the elctro-osmotic mobility (*μ*_0_) of each experiment. **D**. Top: sequence of FUS69-97 with the consensus phosphorylation sites underscored. Bottom: a representative single-molecule time trace of AF647-FUS69-97 after treatment of DNA-dependent protein kinase. Mobility (*μ*) is estimated every 200 ms. Dashed horizontal lines represent the four discrete levels observed in the experiments and are assigned to varying degrees of phosphorylation (0p-3p).

We start by identifying the phosphorylation state of single molecules. We used cAMP-dependent protein kinase catalytical subunit (PKAc) to phosphorylate its classic substrate (kemptide, LRRASLG)^9^. At a low concentration of PKAc (8 nM), the reaction progress can be monitored using Cy5 labeled kemptide and running the stopped reaction at selected time points on an agarose gel (Supplementary Fig. 1). In the gel, the separation of the phosphorylated band from the original band is understood to arise from the added (two) negative charges of the phosphate. We then diluted the stopped reaction to picomolar concentrations and loaded to the ABEL trap. A representative time trace (Fig. 1B) shows single Cy5-kemptide molecules captured for a few seconds each with their diffusion coefficient (*D*) and electrokinetic mobility (*μ*) measured simultaneously. We then pooled all molecules measured at 20 min reaction time (285 molecules) and profiled their diffusivity and mobility on a two-dimensional parameter space (Fig. 1C). The molecules clearly partitioned into two mobility groups with similar diffusivities, bearing resemblance to the two resolved bands on the agarose gel (Supplementary Fig. 1). Conducting similar experiments at different time points further established that the two mobility populations accurately track the phosphorylated and unphosphorylated bands in the gel. In addition, using electrokinetic theory^10^, we calculate the difference in mobility of the unphosphorylated and phosphorylated kemptide to be ∼130 μm/s/V (Supplementary Note 2), similar to the observed difference of 122 μm/s/V. Taken together, we assign the high and low mobility populations to be the unphosphorylated and phosphorylated kemptide, respectively. This assignment uniquely allows us to identify the phosphorylation state in single-molecule traces (Fig. 1B), for example, two molecules at 2 and 14 seconds were unmodified, while the other three molecules were already phosphorylated.

Encouraged by our ability to clearly resolve phosphorylation state from a single serine site, we next challenged the method with a sample where multiple phosphorylation sites are present and could be modified. We chose the 69-97 fragment (FUS69-97) peptide from RNA-binding protein fused in sarcoma (FUS), which is at the heart of the N-terminal low complexity domain^11^. Phosphorylation of the low complexity domain is found to modulate FUS phase separation and disease-associated aggregation^12^. Previous work established that serine/threonine (S/T) followed by glutamine (Q) constitute a phosphorylation site for DNA-dependent protein kinase^12^, therefore two phosphorylation sites are expected for FUS69-97 (Fig. 1D). We labeled FUS69-97 on the N-terminal amine and profiled single peptide’s electrokinetic mobility before and after extended incubation with DNA-dependent protein kinase (Fig. 1D). Strikingly, while unmodified FUS69-97 molecules displayed a high degree of homogeneity (Supplementary Fig. 2), kinase-treated peptides showed four distinct levels of mobility, which we assign to represent molecules with 0, 1, 2, 3 sites phosphorylated. Single-molecule quantification of phosphorylation depth suggests an additional site beyond the consensus (S/T)Q motif and opens up new experimental means to probe phosphorylation biophysics.

We next attempted to watch a single molecule’s phosphorylation and dephosphorylation dynamics in real time. Towards this end we designed a model phosphorylation cycle *in vitro* with PKAc and lambda phosphatase (λPP). Every single substrate molecule is expected to be repeatedly phosphorylated and dephosphorylated, mimicking *in vivo* cycles (Fig. 2A). We first chose PLN20^13^ (cytoplasmic domain of phospholamban, residues 1-20) as the substrate. In bulk, PLN20 can be phosphorylated efficiently by PKAc (Supplementary Fig. 3) with kinetics similar to kemptide (*k*_cat_ ∼ 24 s^-1^, Supplementary Fig. 4) and phosphorylated PLN20 (pPLN20) is a substrate for λPP (Supplementary Fig. 5). When single molecules were monitored without kinase or phosphatase, two static populations of molecules, representing unphosphorylated and phosphorylated PLN20, were observed (Supplementary Fig. 8) similarly to Fig. 1B. With the addition of PKAc, λPP and ATP, we observed dynamic, two-state mobility transitions on single PLN20 molecules (Fig. 2B and Supplementary Fig. 6). These transitions are direct visualizations of charge fluctuations on a single biomolecule due to reversible phosphorylation. Previously, observations of real-time charge fluctuations were limited to microscopic particles^14,15^, here we extend to single biomolecules. The concomitant diffusion coefficient is as high as that of a free peptide (∼150 μm^2^/s) throughout the trace, suggesting that the enzyme bound states (Michalis complex) during catalysis, which would significantly slow down the diffusion, maybe short-lived compared to the time resolution of the measurement (50 ms).

**Figure 2.**
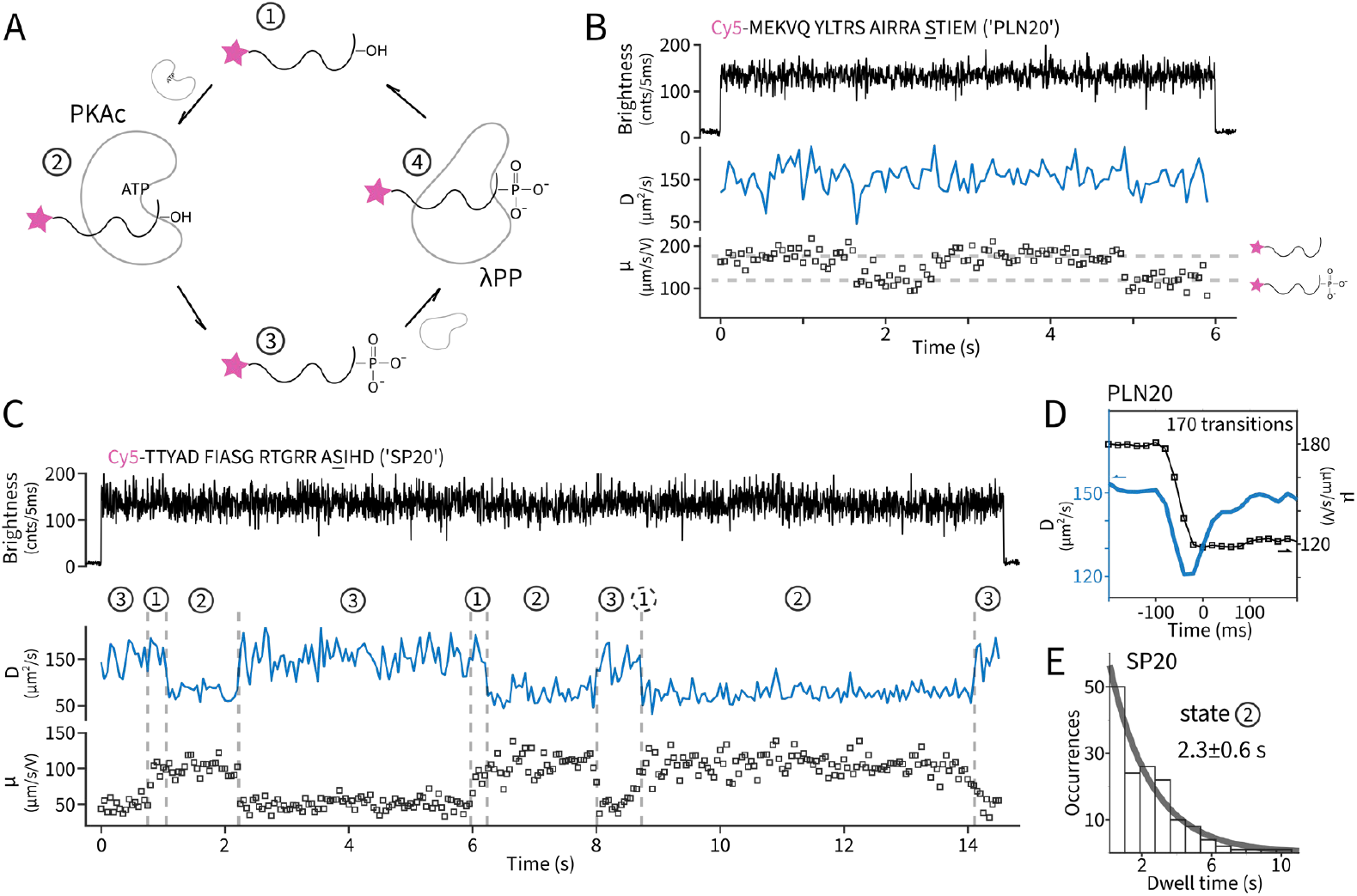
Monitoring dynamic phosphorylation cycles of single molecules. **A.** Scheme of the designed *in vitro* pathway: a single molecule is repeatedly phosphorylated and dephosphorylated by kinase (PKAc) and phosphatase (λPP). **B**. A representative single-molecule time trace of PLN20 as the substrate. The underscored serine in the sequence denotes the phosphorylation site. Diffusivity (*D*) and mobility (*μ*) are estimated every 50 ms. The two horizontal dashed lines represent the mobility levels of unphosphorylated and phosphorylated PLN20. **C**. A representative single-molecule time trace of SP20 as the substrate. Vertical dashed lines represent identified moments of state transitions based on *D* and *μ*. The identified states are numbered according to panel A. Dashed circles denote uncertain state assignments due to limited time resolution (50 ms). **D**. Diffusivity dynamics (blue) near phosphorylation (μ from high to low) transitions (black squares) for PLN20. **E**. Dwell time histogram of state 2 (substrate-PKAc complex) in SP20 experiments. An exponential fit (solid grey) reveals a lifetime of 2.3 ± 0.6 seconds.

We next demonstrate resolving single kinase turnovers. For this we used a “slow” substrate (SP20)^16^ derived from the heat-stable protein kinase inhibitor, inside the PKAc-λPP cycle. Bulk experiments showed SP20 can be phosphorylated by PKAc with a ∼10-fold lower *k*_cat_ (∼2 s^-1^, Supplementary Fig. 4) and a mixture of SP20 and pSP20 can be created with a composition tunable by kinase and phosphatase concentrations (Supplementary Fig. 7). Single-molecule traces of SP20 (Fig. 2C and Supplementary Fig. 8 and 9) displayed similar two-state dynamics in mobility, reflecting the reversible phosphorylation on single substrate molecules, but also with two-state dynamics on molecular diffusivity. In particular, a new *D*∼75 μm^2^/s state now emerged and can last seconds. From control experiments using the corresponding inhibitor peptide (IP20) and structural modeling (Supplementary Fig. 10), we identify this low *D* state to be the peptide-PKAc complex. A closer inspection of the data reveals strict synchrony and order of diffusivity and mobility transitions. For example, high-to-low transitions in *D* always take place when the mobility is high, which we interpret as the ATP-loaded kinase only binding the non-phosphorylated substrates. Meanwhile, high-to-low transitions in mobility are almost strictly in synchrony with low-to-high transitions in *D*, which reflects the kinase’s completion of phosphoryl transfer and releasing the phosphorylated product. Overall, other than the peptide-phosphatase complex (state 4), which are likely too transient^17^ to be observed directly, single-molecules dynamics on SP20 are unambiguously interpreted (Fig. 2C and Supplementary Fig. 9) to follow the state transitions in the counterclockwise direction shown in Fig. 2A. Furthermore, having observed the peptide-PKAc complex in the slow substrate SP20, we went back to the PLN20 data and attempted to probe the transient presence of the kinase bound state. Towards this end, we estimated *D* every 20 ms near identified phosphorylation transitions (*μ* from high to low) and averaged over all such transitions. This analysis revealed a consistent dip of diffusivity that coincide with the mobility switch (Fig. 2D), which is the result of transient PKAc interactions. From the amplitude of the dip (20%), we estimate PKAc stays bound to PLN20 for ∼20 ms.

We have thus demonstrated that electrokinetics expand the measurement dimension of single-molecule fluorescence experiments to include phosphorylation and in general, charge-related modifications. Our implementation is on the ABEL trap, but other experimental platforms are possible^18–20^. Direct phosphorylation sensing opens many future directions in studying structural, catalytic and functional processes driven by kinases and phosphatases with single-molecule resolution. Here we provide an example of identifying the rate-limiting step in SP20 phosphorylation by PKAc. It was observed directly that SP20 forms a stable complex with PKAc, before being released as the phosphorylated product (Fig. 2C). The dwell times of the complex (state 2) are exponentially distributed with a mean of 2.3 ± 0.6 seconds (Fig. 2D). We hypothesized that the complex represents the product (pSP20) bound to PKAc after phosphoryl transfer. Indeed, when we monitored single-molecule dynamics using phosphorylated SP20, PKAc and ADP to mimic the post catalysis condition, similar long dwells (2.1 ± 0.7 s) of the peptide-PKAc complexes were observed (Supplementary Fig. 11). This observation suggests that product trapping, not phosphoryl transfer, is rate-limiting and ADP release likely happens after product release, consistent with recent biochemical evidence^16^. Ultimately, we envision that by combining high-resolution single-molecule Förster resonance energy transfer (smFRET)^21^, detailed structural insights during or after phosphorylation may be obtained.

## Online Methods

### Enzymes and peptide substrates

cAMP-dependent Protein Kinase A catalytic subunit (PKAc, 13 μM) and Lambda Protein Phosphatase (λPP, 20 μM) were purchased from New England Biolabs (Ipswich, MA) and used without further purification. Peptides (kemptide, PLN20, SP20), both labeled and unlabeled, were custom synthesized by Anaspec (Supplementary Table 1). FUS69-97 peptide was a gift from Robert Tycko and labeled with AF647-NHS ester (Lumiprobe) on the N-terminal amine following manufacturer’s recommended procedure and purified using reverse phase chromatography.

### Phosphorylation reactions

Kemptide phosphorylation reactions were conducted using 60 μM Cy5-kemptide, 8 nM PKAc and 1 mM ATP in PKAc reaction buffer (50 mM Tris-HCl pH 7.5, 10 mM MgCl_2_, 0.1 mM EDTA, 2 mM DTT, 0.01% Brij 35). Reactions were stopped at selected times by adding 50 mM EDTA and incubation at 70 °C for 10 min. The stopped reactions were diluted to 10 pM Cy5-kemptide in a buffer containing 20 mM HEPES pH7.5, 2 mM Trolox, 10 mM NaCl for single-molecule trapping experiments. Dynamic phosphorylation and dephosphorylation reactions were prepared with 10 pM Cy5-PLN20 or Cy5-SP20, 0.26 μM PKAc, 0.4 μM λPP in a buffer containing 20 mM HEPES pH 7.2, 2 mM Trolox, 100 mM NaCl, 5 mM MgCl_2_ 0.4 mM MnCl_2_, 0.4 mM ATP. Phosphorylation of FUS69-97 fragment was achieved by DNA-dependent protein kinase (DNA-PK). The reaction was carried out in DNA-PK kinase reaction buffer (40 mM Tris-HCl, pH 7.5, 20 mM MgCl_2_, 0.1 mg/ml BSA) with 10 μg/mL calf thymus DNA, 2.6 μM AF647-FUS69-97, 0.9 u/uL DNA-PK kinase (Promega V4106), 0.2 mM ATP and incubated at room temperature overnight. The reaction was diluted to 10 pM of AF647-FUS69-97 in a buffer containing 20 mM HEPES pH7.7, 2 mM Trolox and 20 mM NaCl for single-molecule trapping experiments.

### ABEL trap implementation

The ABEL trap with acousto-optic beam scanning, 633 nm excitation and a focus lock was implemented based on a published design on a custom microscope base (MadCity Labs RM21)^22,23^. Briefly, a single-molecule’s motion is monitored photon-by-photon within a small trapping region (∼3 μm radius circle) in solution by rapid beam scanning and a Kalman filter. Feedback voltages were applied accordingly with minimum delays to move the molecule towards the center of the trapping region. As a result, a single molecule is controlled to diffuse within the circular trapping region and stays within the detection volume of a confocal fluorescence microscope for extended times. Typical capture durations in this work on single peptides were 1-20 seconds, limited by photobleaching. Termination of a single-molecule trapping event frees up the trap for another molecule to randomly enter and be captured. Molecules are thus measured sequentially with interspersed background regions, exemplified by Fig. 1B and 1D.

### ABEL trap experiments

Before sample loading, fused silica sample chambers were cleaned, coated with 2 mg/ml polyethyleneimine (in 20 mM HEPES pH 7, 150 mM NaCl), rinsed and followed by coating with 2 mg/ml polyacrylic acid. The sample was supplemented with 100 nM protocatechuate 3,4-dioxygenase (OYC America, in house purified by size exclusion chromatography) and 2 mM protocatechuic acid (Sigma) for enzymatic oxygen removal before loading. Single-molecule trapping experiments were conducted at 23 °C and typically run for 0.5-2 hours till ∼200-500 molecules were collected.

### Data analysis

All data analysis was performed with customized software written in Matlab. The electrokinetic mobility (*μ*) and diffusion coefficient (*D*) of a trapped single molecule were estimated by a maximum likelihood framework developed previously^7^. Briefly, the in-trap motion is tracked photon-by-photon and decomposed into a voltage-dependent, electrokinetic response (for estimating *μ*) and a voltage-independent, diffusive component (for estimating *D*) by an expectation-maximization algorithm. For time-dependent estimations, the algorithm runs on non-overlapping time windows of 50 ms-200 ms. Synchronized dynamics of *D* and *μ* were analyzed using a joint change point finding algorithm^24^.

## Acknowledgement

We thank Rob Tycko for the gift of FUS69-97 peptide, Miles Lee for electrokinetic mobility calculations, Elif Karasu for assistance with sample preparation and Bok-Eum Choi for in house purification of protocatechuate 3,4-dioxygenase. This work was supported by the Intramural Research Program of the National Institute of Diabetes and Digestive and Kidney Diseases, NIH.

