## Supplementary material for "Electrokinetic detection of single-molecule phosphorylation": SI

Quan Wang

Laboratory of Chemical Physics, National Institute of Diabetes and Digestive and Kidney Diseases, National Institutes of Health, Bethesda, Maryland 20892-0520, United States

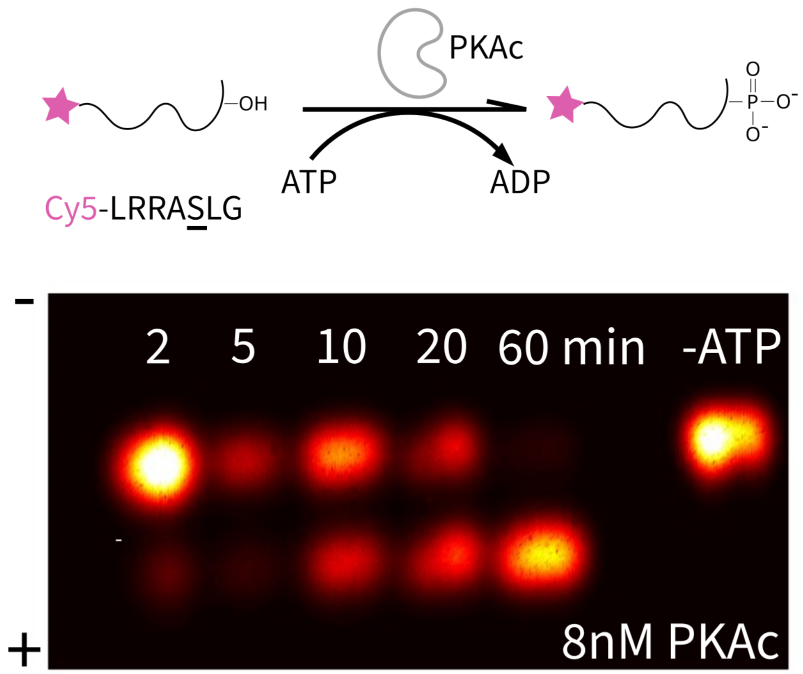

**Supplementary Figure 1. Monitoring phosphorylation reaction progress on agarose gel.** Top: reaction scheme and the sequence of kemptide. The underscored serine residue designates the phosphorylation site. The reaction was conducted using 60 µM Cy5-kemptide, 8 nM PKAc, 1 mM ATP, PKAc reaction buffer (50 mM Tris-HCl pH 7.5, 10 mM $MgCl_2$, 0.1 mM EDTA, 2 mM DTT, 0.01% Brij 35). In the negative control reaction, ATP was not added. At selected times, the reaction was sampled and stopped by adding 50 mM EDTA and heating to 70 °C for 10 min. Bottom: fluorescence gel image of the stopped reactions at different times and the control reaction without ATP. Each stopped reaction was diluted to ~50 nM (peptide) and ran on 0.8% agarose gel at 100V for 30 min. The gel was imaged on a home-built scanner using 642 nm excitation. The polarity of the applied voltage was shown on the left.

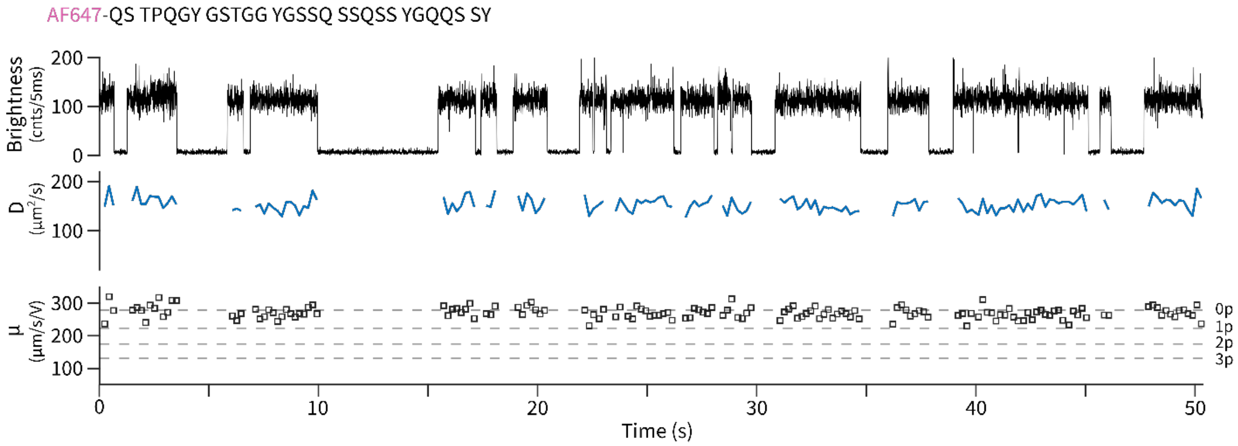

**Supplementary Figure 2. Unphosphorylated FUS69-97 peptide molecules show uniformly high values of electrokinetic mobility (μ)**. A representative ABEL trap trace of AF647 labeled FUS69-97, with brightness, diffusion coefficient (*D*) and electrokinetic mobility ($\mu$). The dashed lines represent the four levels of $\mu$ observed in the experiment shown in Fig. 1D of the main text. To conduct this experiment, AF647 labeled FUS69-97 was diluted to 10 pM in the trapping buffer containing 20 mM HEPES pH7.7, 2 mM Trolox and 20 mM NaCl, with an oxygen scavenging system (100 nM PCO, 2 mM PCA). Diffusion coefficient (*D*) and electrokinetic mobility ($\mu$) were estimated every 200 ms.

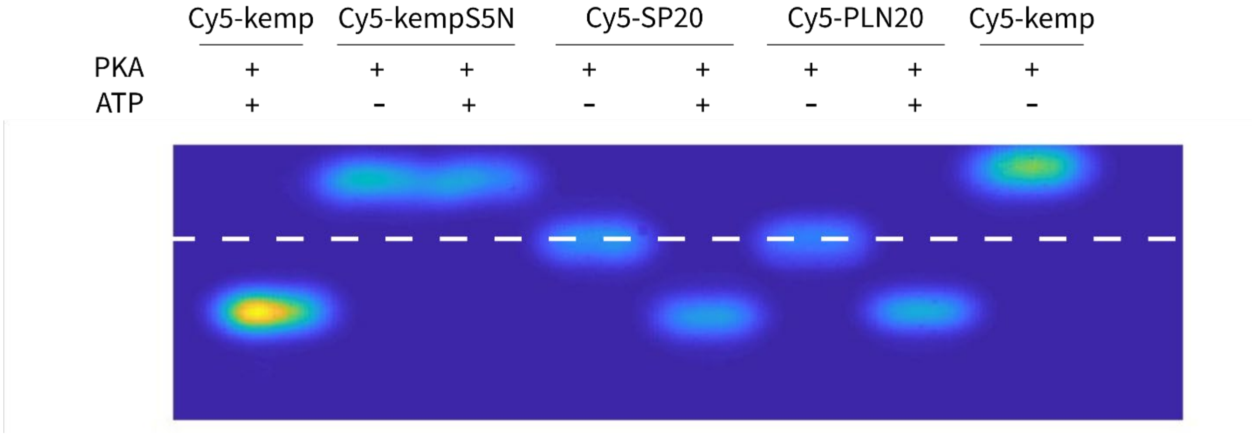

**Supplementary Figure 3. Phosphorylation of peptide substrates by PKAc.** Fluorescence gel image of experiments that test if selected peptide substrates can be phosphorylated by PKAc. The dashed line indicates the location of the loading wells. Each reaction contains 70 nM Cy5-labeld peptide,100 nM PKAc with or without ATP (200 µM) in PKAc reaction buffer (50 mM Tris-HCl pH 7.5, 10 mM $MgCl_2$, 0.1 mM EDTA, 2 mM DTT, 0.01% Brij 35). The reactions were allowed to proceed for 4 hours and stopped by adding 50 mM EDTA and heating to 95 °C for 10 min. The stopped reactions were resolved in 0.8% agarose gel and imaged.

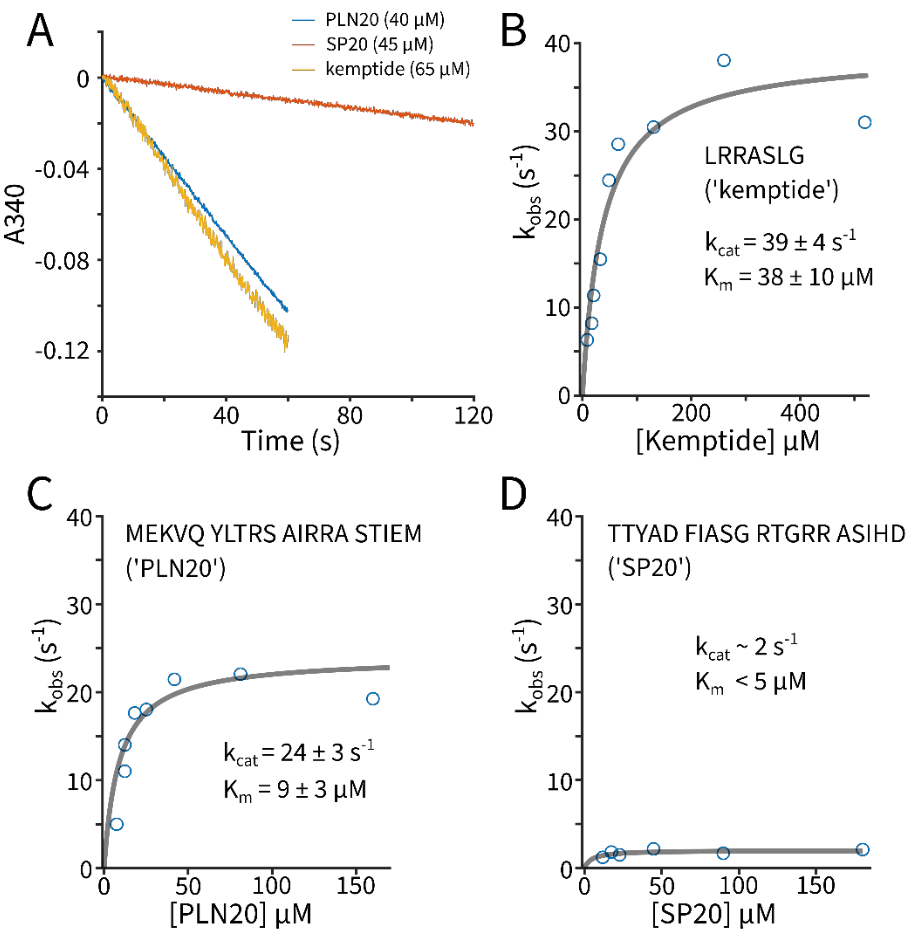

**Supplementary Figure 4. Steady-state phosphorylation kinetics of PKAc on peptide substrates. A**) The loss of 340 nm absorption as a function of time for the three peptide substrates used in this work. Kinase activity was measured by coupling the generation of ADP with the oxidation of NADH by pyruvate kinase and lactate dehydrogenase[1]. Each reaction contains 0.6 mg/ml NADH, 3 mM phosphoenolpyruvate, 0.02 u/uL L-lactate dehydrogenase, 0.02 u/uL pyruvate kinase, 0.1 mg/ml BSA, 1 mM ATP, 13 nM PKAc in a buffer containing 20 mM HEPES pH7.5, 20 mM NaCl, 1 mM $MgCl_2$. Reactions were initiated by adding peptide substrates and absorption at 340 nm were recorded as a function of time, up to 120 seconds. Michalis-Menten plots of substrate **B**) kemptide, **C**) PLN20 and **D**) SP20. Measured rates ($k_{obs}$) were fitted (solid gray lines) using $k_{cat} \times c/(c + K_m)$. Errors represent 68% confidence intervals. For SP20, the gray curve is a direct plot of the above equation with $k_{cat} = 2\ s^{-1}$ and $K_m = 5\ \mu M$.

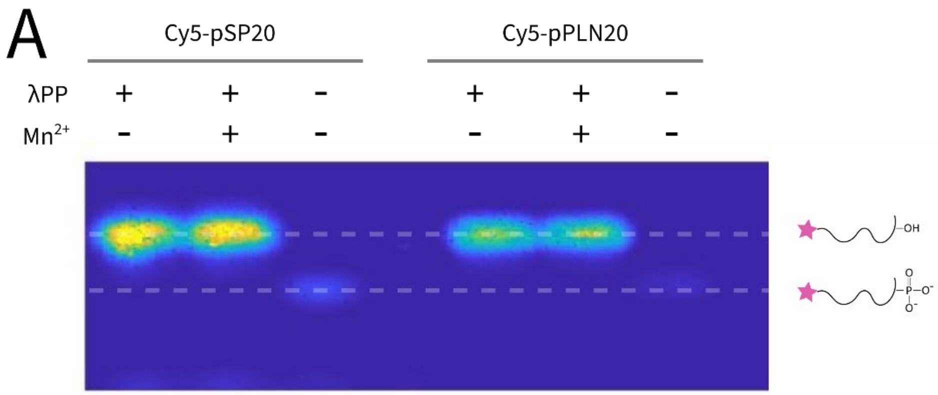

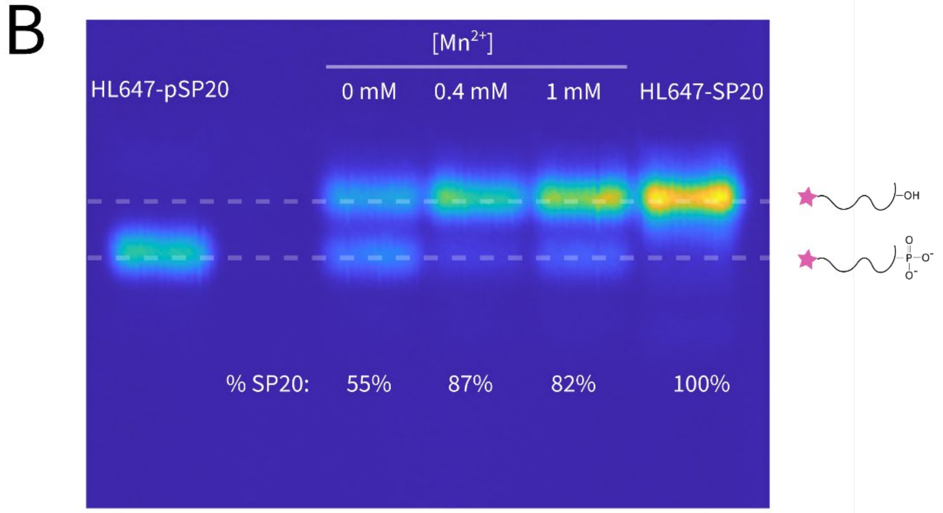

**Supplementary Figure 5. Lambda phosphatase (λPP) dephosphorylates pPLN20 and pSP20.** **A**) Fluoresce gel image of bulk experiments that test whether λPP recognizes pSP20 and pPLN20 as substrates. In these experiments, 0.75 µM of phosphorylated peptides (pPLN20 and pSP20) were incubated with 0.4 µM λPP in a buffer containing 50 mM HEPES pH7.5, 100 mM NaCl, 2 mM DTT, 0.01% Brij 35. In reactions that contains $Mn^{2+}$, 1 mM $MnCl_2$ was added to the reaction mix. Reactions were allowed to proceed for 1 hour and subsequently diluted 10× in 50 mM Tris-HCl buffer pH8 and ran on 1% Agarose gel (100 V, 30 min). The gel was imaged on a home-built imager using 642 nm excitation and a Cy5 emission filter set. **B**) Fluorescence gel image of bulk experiments that test $Mn^{2+}$ dependence of λPP activity. Stock λPP was diluted in buffers containing 20 mM HEPES pH7.2, 2 mM Trolox, 20 mM NaCl, 5 mM $MgCl_2$ with three different concentrations of $MnCl_2$ (0, 0.4 mM or 1 mM). After incubation for 100 min, 10 µM of HL647 labeled pSP20 was added and the reactions were allowed to proceed for 1 min before quenching. The degree of dephosphorylation was assayed by running the quenched reactions on 1% agarose gel. The results show that λPP loses activity in a buffer without $Mn^{2+}$ and justify the inclusion of 0.4 mM $Mn^{2+}$ in the phosphorylation cycle experiments.

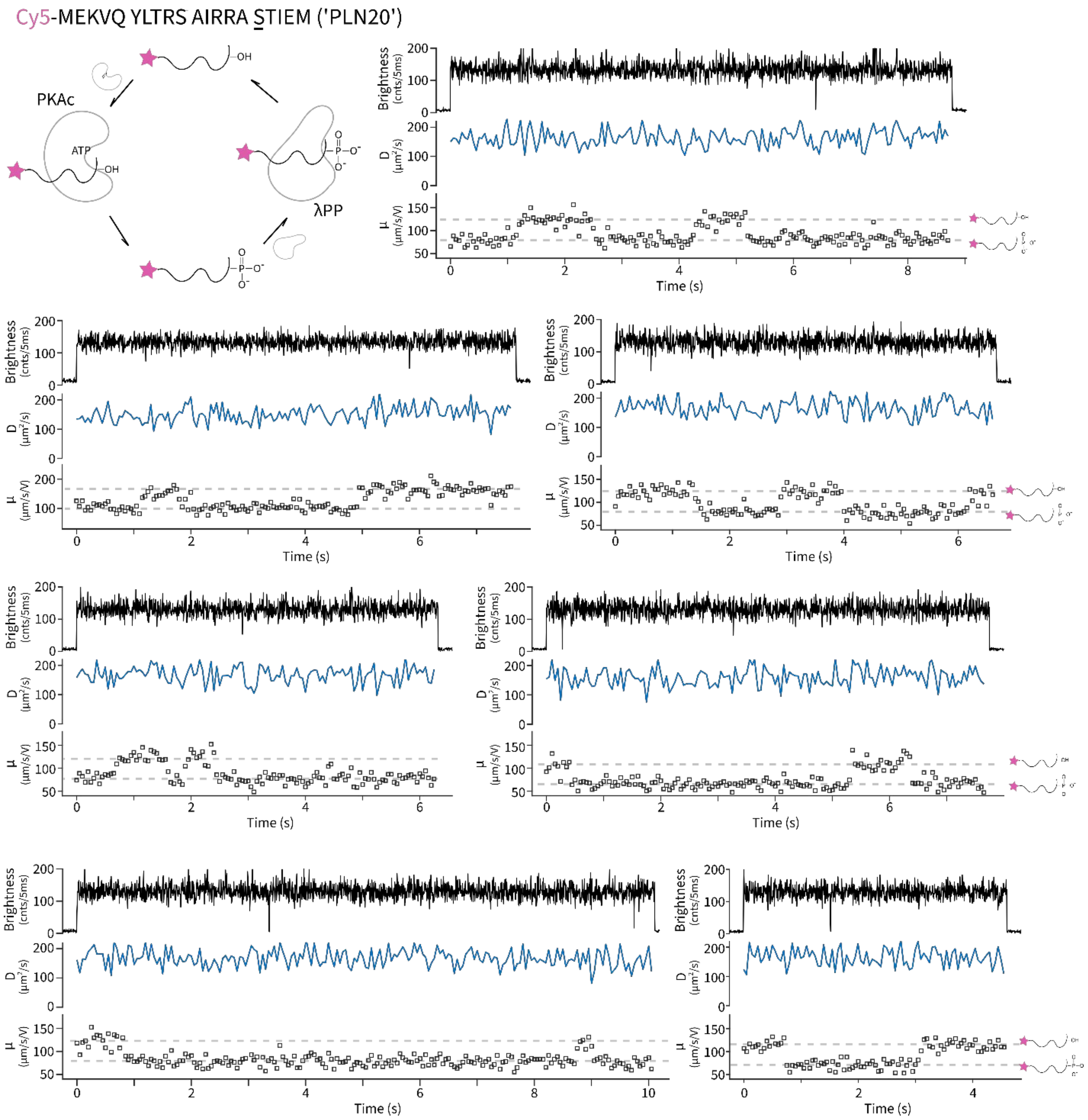

**Supplementary Figure 6. Additional single-molecule traces of PLN20 undergoing phosphorylation and dephosphorylation cycles.** See also Fig. 2B of the main text.

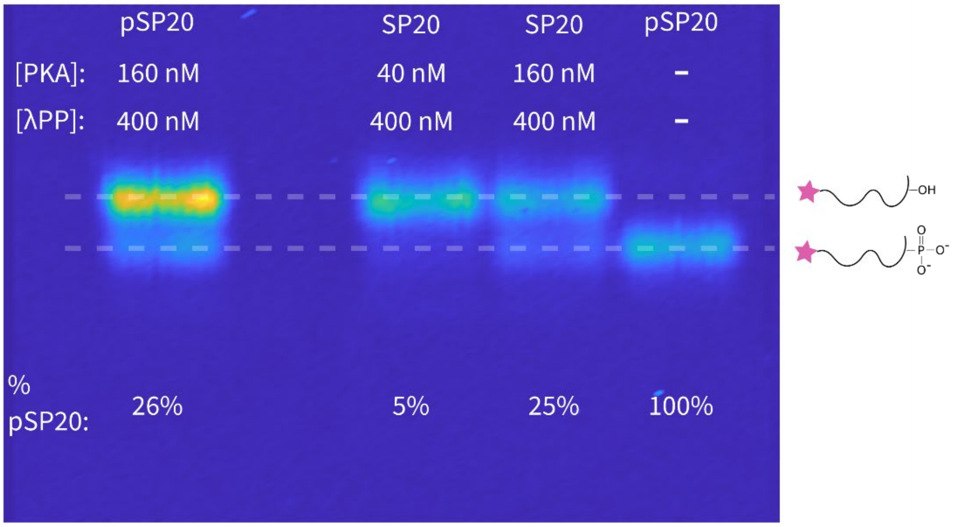

**Supplementary Figure 7. PKAc and λPP together creates steady-state mixtures of unphosphorylated and phosphorylated SP20 at the ensemble level.** Fluorescence gel image of bulk experiments that validates the phosphorylation and dephosphorylation cycle (Fig. 2A of main text). Each reaction contains 5 nM of HL647 labeled SP20 (or pSP20), PKAc, λPP, 0.4 mM $MnCl_2$, 0.4 mM ATP in 20 mM HEPES pH7.2, 2 mM Trolox, 20 mM NaCl and 5 mM $MgCl_2$. After 10 min incubation at room temperature, the reactions were quenched by 10% acetic acid, 50 mM EDTA and heating to 70 °C. The reaction products were separated on 1% agarose gel (100 V, 30 min) and quantified by the fluorescence intensity of the bands. The steady-state fraction of pSP20 is independent of the initial starting peptide (SP20 or pSP20) and depends on the concentration of PKAc and λPP.

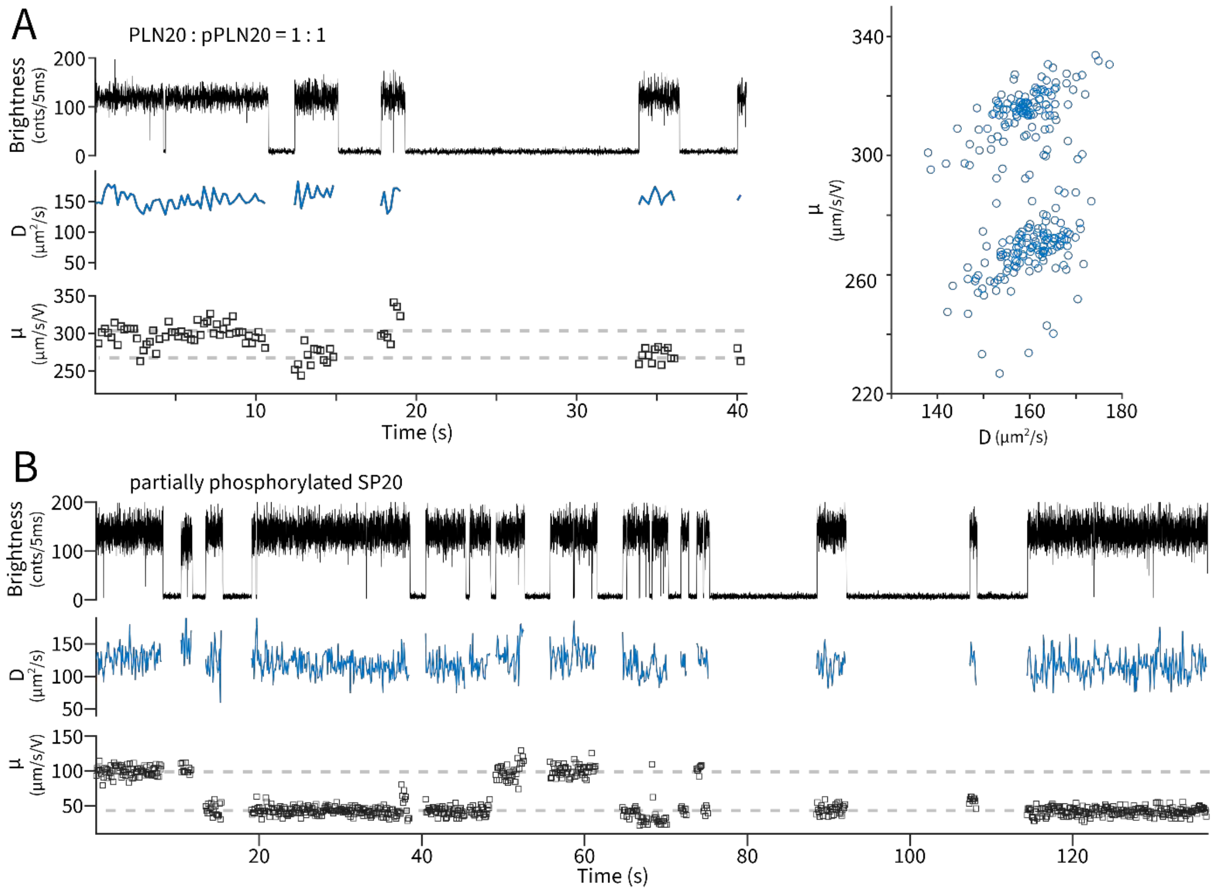

**Supplementary Figure 8. No single-molecule dynamics on static mixtures of both phosphorylated and unphosphorylated peptide substrates. A)** Left: a representative single-molecule trace of a 50:50 mixture composed of PLN20 and pPLN20. Diffusivity and mobility are calculated with 200 ms time resolution. Right: Diffusivity-mobility profile of 347 measured molecules. The cluster with lower mobility is pPLN20. Phosphorylated PLN20 (pPLN20) was generated enzymatically. Trapping experiments were conducted in 20 mM HEPES pH7.5, 2 mM Trolox, 20 mM NaCl, 50 nM PCO and 2 mM PCA. **B)** a representative single-molecule trace of partially phosphorylated SP20. To generate a mixture of SP20 and pSP20, 70 nM of SP20 was incubated with 10 nM PKAc, 200 µM ATP in PKAc reaction buffer (50 mM Tris-HCl pH 7.5, 10 mM $MgCl_2$, 0.1 mM EDTA, 2 mM DTT, 0.01% Brij 35). The reaction was allowed to proceed for 10 seconds before quenched with 20 mM EDTA and heating at 70 ºC for 20 min. The stopped reaction was diluted to 20 pM (peptide) in 20 mM HEPES pH7.5, 2 mM Trolox, 10 mM NaCl, 1 mM $MgCl_2$ and 10% glycerol. Oxygen was removed in the final solution using 50 nM PCO and 2 mM PCA. Diffusivity and mobility are calculated with 50 ms time resolution. Single peptide molecules show either $\mu$ ~ 100 µm/s/V (SP20) or $\mu$ ~ 50 µm/s/V (pSP20) with no discernible dynamics.

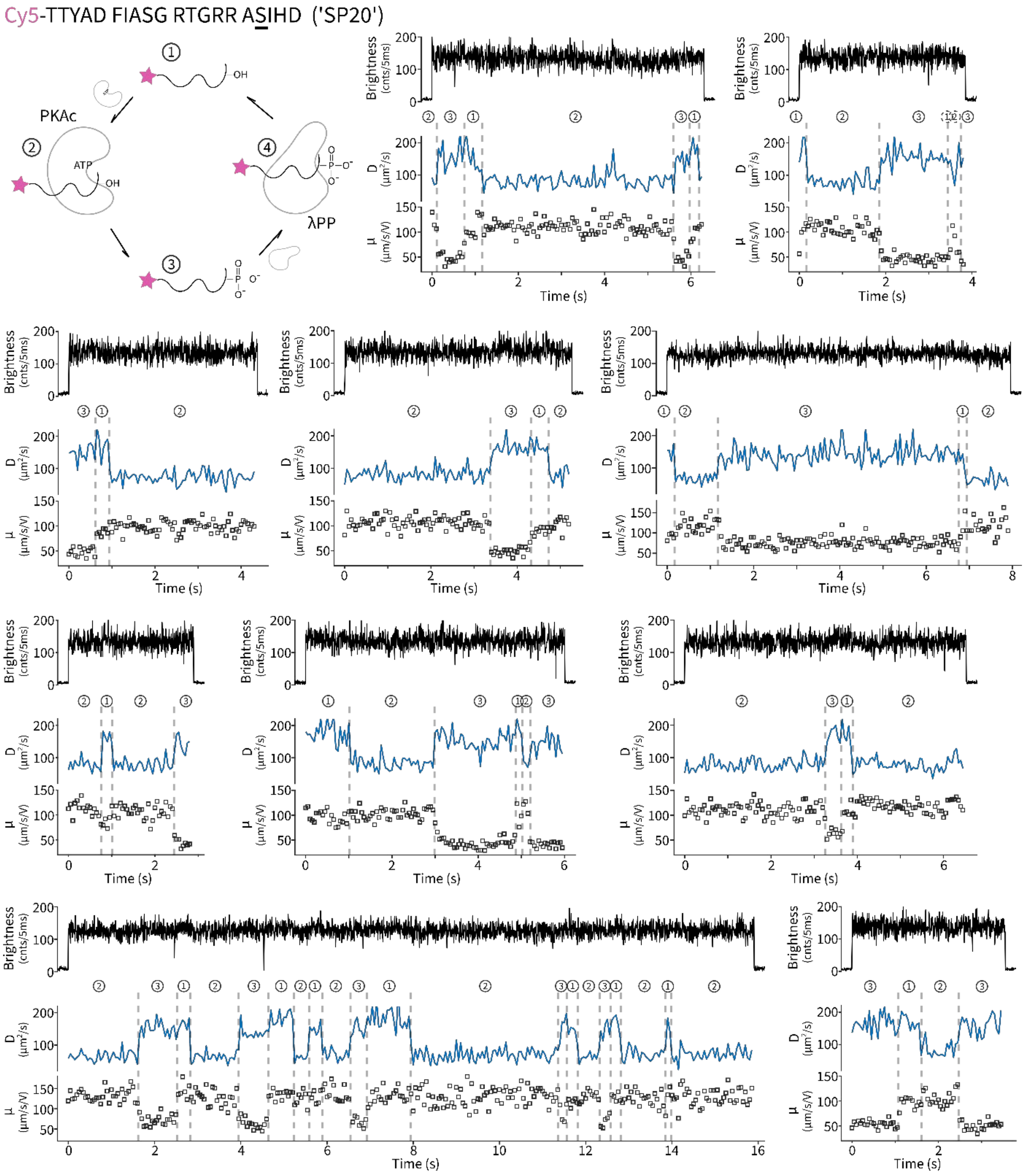

**Supplementary Figure 9. Additional single-molecule traces of SP20 undergoing phosphorylation and dephosphorylation cycles.** See also Fig. 2C of the main text.

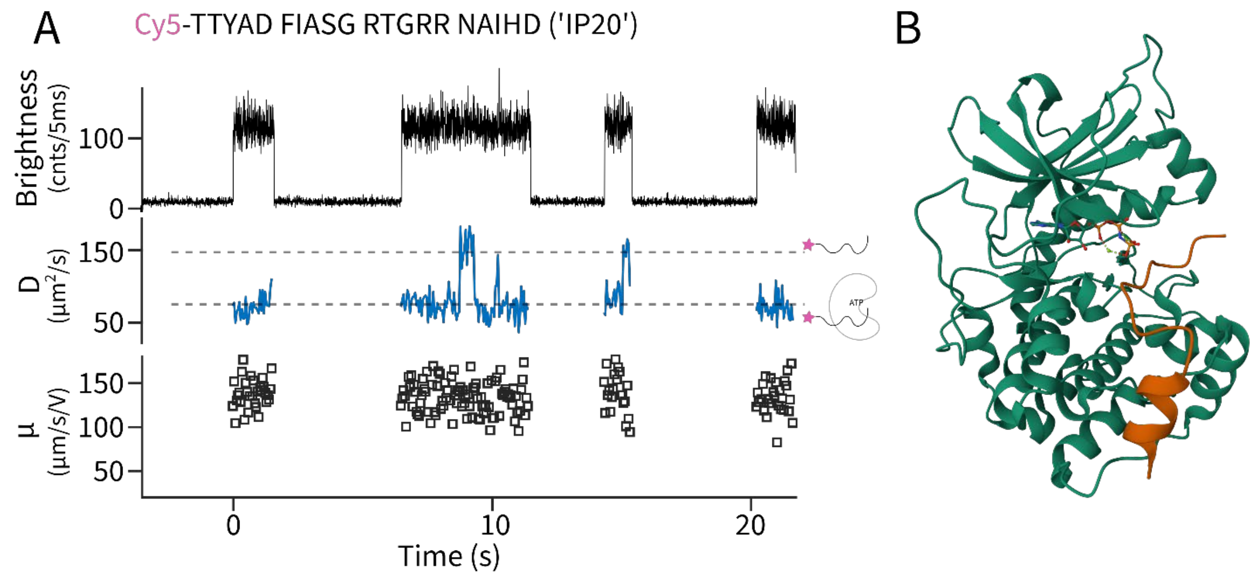

**Supplementary Figure 10. Probing the peptide-PKAc complex using IP20 and structural modeling. A)** A representative ABEL trap time trace of Cy5-IP20 in the presence of PKAc and ATP. Diffusivity and mobility are estimated every 50 ms. The low diffusivity state ($D \sim 75$ µm$^2$/s) is identified to be the IP20-PKAc complex. Transient unbinding and rebinding (e.g. at ~9.5s) are frequently observed. In this experiment, Cy5 labeled IP20 was diluted to 10 pM with 260 nM PKAc, 0.4 mM ATP, in the trapping buffer that contains 20 mM HEPES pH7.5, 2 mM Trolox, 100 mM NaCl, 5 mM $MgCl_2$ and an oxygen scavenger system (100 nM PCO with 2 mM PCA). **B)** Crystal structure of the IP20-PKAc complex (pdb:4DH8) with AMPPNP as the ATP analog: PKAc is colored in green, IP20 is colored in orange and AMPPNP is rendered as ball and sticks. From this structure, hydrodynamic modeling (HydroPro[2]) predicts a diffusion coefficient of 74.8 µm$^2$/s, supporting the assignment of the low $D$ state in experiments as the peptide-PKAc complex.

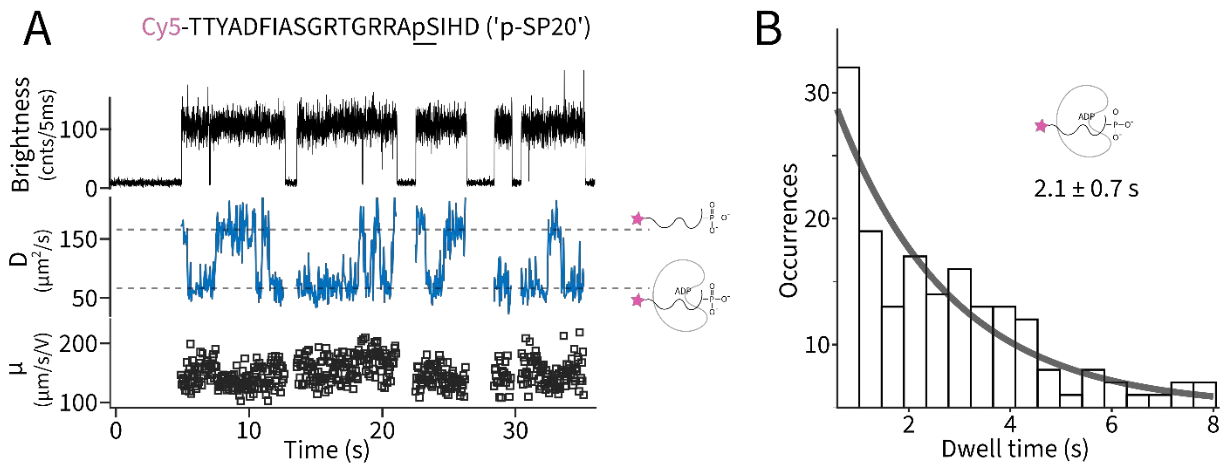

**Supplementary Figure 11. Probing single molecules of the kinase-product complex. A**) A representative single-molecule trace of Cy5-pSP20 (10 pM) captured in the presence of 260 nM PKAc, 1 mM ADP in a buffer containing 20 mM HEPES pH 7.2, 2 mM Trolox, 100 mM NaCl, 5 mM $MgCl_2$, 0.4 mM $MnCl_2$. In this experiment, diffusion coefficient was observed to transition between a $D$ ~ 150 $\mu m^2/s$ state and a $D$ ~ 75 $\mu m^2/s$ state, which we interpret to represent the binding/unbinding dynamics of pSP20 with ADP loaded PKAc. Note that slight mobility differences between the free and PKAc-bound peptides could also be resolved. **B**) Dwell time histogram of the peptide-PKAc complex with a single exponential fit (solid gray) extracts the lifetime to be 2.1 ± 0.7 seconds.

**Supplementary Table S1. Peptide sequences**

| Name | Sequence (N→C) | Nominal charge (pH 7) |
|---|---|---|
| kemptide | LRRASLG-$NH_2$ | +3 |
| Cy5-kemptide | Cy5-LRRASLG-$NH_2$ | +1 |
| Cy5-kemptideS5N | Cy5-LRRANLG-$NH_2$ | +1 |
| Cy5-IP20 | Cy5-TTYADFIASGRTGRRNAIHD-$NH_2$ | 0 |
| PLN20 | MEKVQYLTRSAIRRASTIEM | +2 |
| Cy5-PLN20 | Cy5-MEKVQYLTRSAIRRASTIEM | 0 |
| SP20 | TTYADFIASGRTGRRASIHD-$NH_2$ | +2 |
| Cy5-SP20 | Cy5-TTYADFIASGRTGRRASIHD-$NH_2$ | 0 |
| HL647-SP20 | HiLyte647- TTYADFIASGRTGRRASIHD | |
| AF647-FUS69-97 | AF647-QS TPQGY GSTGG YGSSQ SSQSS YGQQS SY | -4 |

Red: positively charged residues

Green: negatively charged moieties (amino acids and dyes)

Underscored: consensus phosphorylation sites

-$NH_2$: C-terminal amidation.

**Supplementary Note 1 Electroosmotic and electrophoretic mobilities**

In all ABEL trap experiments conducted in this work, we modified the microfluidic chips with polyacrylic acid to introduce negatively charged surfaces at neutral pH. This surface modification prevents non-specific adsorption of proteins[3] and more importantly, introduces strong electroosmotic flows[4] that maintain trapping of single molecules undergoing charge reversals (e.g. from +1 charge to -1 charge). We briefly outline the principles below.

The overall electrokinetic response of a single molecule in a microfluidic channel under applied voltage is made up of two components: electrophoresis (EP) and electroosmosis (EO). Electrophoresis refers to the direct motion of charged object under the electric field (i.e. towards the anode for a negatively charged molecule). Electroosmotic flow, on the other hand, is generated by charged surfaces via a counterion gradient (Debye layer). Given that the surface charges are fixed and the counterions are mobile, the result is a bulk flow with a direction determined by the type of the counterion (i.e. towards the cathode for negatively charged surfaces). We thus have $\mu = \mu_{EO} - \mu_{EP}$ with positive $\mu$ pointing to the cathode. If we make $|\mu_{EO}| > |\mu_{EP}|$, which can be achieved by increasing the charge density on the surfaces, charge reversal ($\mu_{EP}$ flips sign) would not change the sign of $\mu$. In other words, electroosmotic mobility $\mu_{EO}$ adds a constant offset to the overall mobility response.

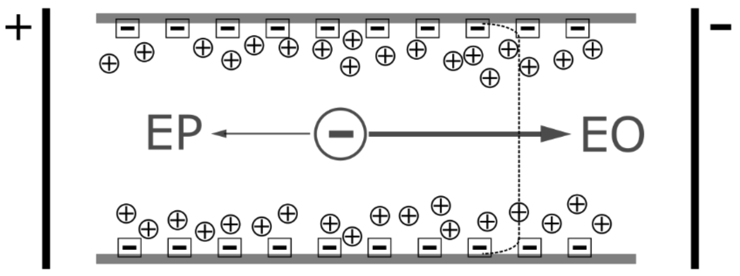

Figure S12 **Electroosmotic and electrophoretic flows with negatively charged channel walls.** The top and bottom channel walls are drawn in grey solid lines. Fixed charges are in squares and mobile counterions are in circles. The particle of interest, assumed to be negatively charged, is drawn in the center. Directions of electrophoretic migration (EP) and electroosmotic flow (EO) are drawn as arrows, in response to the external field as shown.

**Supplementary Note 2 Estimation of the mobility difference between phosphorylated and unphosphorylated kemptide**

In classical electrokinetic theory[5], the electrophoretic mobility of a charged particle in buffered aqueous solution can be computed using Henry's equation,

$$\mu_E = \frac{2\varepsilon\zeta}{3\eta} f_1(\kappa a)$$

where $\eta$ is the dynamic viscosity of water, $\varepsilon$ is the relative permittivity, $\zeta$ is the zeta potential. For a sphere with charge $Z$ and radius $a$, the zeta potential can be estimated using

$$\zeta = \frac{eZ}{4\pi\varepsilon a(1+\kappa a)}$$

where $\kappa$ is the ionic strength,

$$\kappa = \sqrt{\frac{e^2 \Sigma n_i z_i^2}{\varepsilon kT}}$$

Particle radius $a$ can be estimated using the simultaneously measured diffusion coefficient $D_t$ using the Stokes–Einstein relation

$$a = \frac{kT}{6\pi\eta D_t}$$

The function $f_1(\kappa a)$ depends on particle shape. Approximate formula exists to calculate $f_1$ for arbitrary values of $\kappa a$. For $0 < \kappa a < 2$, $f_1(\kappa a) \approx 1$ within 5% error. For kemptide in 10 mM NaCl, debye length $\kappa^{-1}$ = 0.31 nm. Using a measured diffusivity of $D_t$ = 230 μm$^2$/s, $a$ = 0.95 nm. After phosphorylation, $Z$ changes from +1 to -1. We approximate that applied voltage on the microfluidic chip creates a uniform electric field over a distance of 100 μm (Ref [6]). Plugging these numbers to the Henry's equation, we calculate $\mu_E$ for kemptide to be ~65 μm/s/V and the mobility difference between unmodified and phosphorylated kemptide to be 130 μm/s/V.
